# Landscape of semi-extractable RNAs across five human cell lines

**DOI:** 10.1101/2022.10.03.510572

**Authors:** Chao Zeng, Takeshi Chujo, Tetsuro Hirose, Michiaki Hamada

## Abstract

Phase-separated membraneless organelles often contain RNAs that exhibit unusual semi-extractability upon the conventional RNA extraction method, and can be efficiently retrieved by needle shearing or heating during RNA extraction. Semi-extractable RNAs are promising resources for understanding RNA-centric phase separation. However, limited assessments have been performed to systematically identify and characterize semi-extractable RNAs. In this study, 1,074 semi-extractable RNAs were identified across five human cell lines, including ASAP1, DANT2, EXT1, FTX, IGF1R, LIMS1, NEAT1, PHF21A, PVT1, SCMH1, STRG.3024.1, TBL1X, TCF7L2, TVP23C-CDRT4, UBE2E2, ZCCHC7, ZFAND3, and ZSWIM6, which exhibited consistent semi-extractability. By integrating publicly available datasets, we found that semi-extractable RNAs tend to be distributed in the nuclear compartments but are dissociated from the chromatin. Long and repeat-containing semi-extractable RNAs act as hubs to provide global RNA-RNA interactions. Semi-extractable RNAs were divided into four groups based on their k-mer content. The NEAT1 group preferred to interact with paraspeckle proteins, such as FUS and NONO, implying that RNAs in this group are potential candidates of architectural RNAs that constitute nuclear bodies.

## INTRODUCTION

Liquid-liquid phase separation (LLPS) is a biological phenomenon in which macromolecules, such as proteins or nucleic acids, are spatially organized into membrane-less organelles (also called biomolecular condensates) (1). Membrane-less organelles (MLOs) usually maintain their stable structures through multivalent interactions of molecules that act in diverse biological processes ranging from macromolecular biogenesis to gene regulation (2, 3, 4). MLOs are highly dynamic structures, whose components rapidly exchange between other condensates and the surrounding milieu (5, 6, 7, 8, 9), implying that MLOs are sensitive to internal and external signals. LLPS provides a new framework for our understanding of human health and disease (10, 11, 12). Phase-separated MLOs that have been discovered and studied include the nucleolus, paraspeckle, nuclear speckle, Cajal body, PML nuclear body, P-body, stress granule, germ granule, and mRNP granule (3). The role of proteins in LLPS and their regulation has been the focus of attention (1, 13, 14, 15). However, based on accumulating evidence, RNAs, especially long noncoding RNAs (lncRNAs), play a crucial role in the process of phase separation (16, 17, 18, 19, 20, 21).

As a remarkable example, nuclear paraspeckle assembly transcript 1 (NEAT1) is an architectural lncRNA that mediates the assembly of paraspeckles by driving phase separation (22, 23, 24, 25). Two major isoforms are generated from the NEAT1 gene locus, and the longer isoform NEAT1 2 serves as a molecular scaffold for the formation of RNA-protein and RNA-RNA interactions (19, 26). Paraspeckles form a core-shell spheroidal structure, in which the shell contains the 5^*′*^ and 3^*′*^ regions of NEAT1 2 and some specific proteins, whereas the core consists of the middle region of NEAT1 2 and Drosophila behaviour/human splicing (DBHS) proteins (27). According to further studies, the NEAT1 2 middle region contains redundant subdomains that sequester RNA-binding proteins (RBPs), such as non-POU domain-containing octamer-binding protein (NONO) and splicing factor proline and glutamine rich (SFPQ), to initiate paraspeckle assembly (28). Note that both NONO and SFPQ are members of the DBHS family of proteins. Interestingly, when a conventional RNA extraction method using AGPC (acid guanidinium thiocyanate-phenol-chloroform) reagent such as TRIzol is employed, most of the NEAT1 is retained in the protein layer between the aqueous phase and organic phase, resulting in a low extraction level. However, after the phase-separated structures are disrupted by an improved RNA extraction through needle shearing or heating, NEAT1 is released into the aqueous solution, and its extraction level can be 20-fold higher than that obtained via the conventional method. Such property of NEAT1 is termed as ‘semi-extractability’ (29). The semi-extractability of NEAT1 strongly depended on the prion-like domain of a paraspeckle RBP, FUS, implying that extensive multivalent interactions may cause semi-extractability (22). In addition to NEAT1, several other newly detected semi-extractable RNAs were observed to form granule-like foci in a previous study (29). Accordingly, RNAs in the phase-separated structures may commonly possess semi-extractability owing to multivalent forces. The systematic identification and characterization of semi-extractable RNAs could aid in the discovery of RNAs associated with phase separated MLOs and provide insights into LLPS biology.

In this study, we developed a bioinformatic pipeline to define 1,074 semi-extractable RNAs for the first time in five human cell lines. As controls, we also identified 6,695 extractable RNAs from all five cell lines. Extractable RNAs were defined as RNAs without pronounced expression changes using the improved RNA extraction method, indicating that they were recovered well by the conventional method. Compared with extractable RNAs, semi-extractable RNAs prefer to be transcribed from repressed and repetitive/heterochromatin regions that are clustered in the nuclear compartments. Long and AU-rich semi-extractable RNAs contain more repetitive sequences than expected and interact frequently with other RNAs. Semi-extractable RNAs can be broadly classified into four different groups based on their sequence composition, with the semi-extractable RNAs of the NEAT1 group preferring to bind paraspeckle RBPs (e.g., NONO and FUS), suggesting their potential role as architectural RNAs.

## MATERIALS AND METHODS

### Cellular RNA extraction and sequencing

A10, A549, HEK293 and HeLa cells were grown in DMEM (GIBCO) supplemented with 10% fetal bovine serum (FBS) in a humidified, 37°C atmosphere with 5% CO_2_. HAP1 cells were grown similarly in IMDM (GIBCO) supplemented with 10% FBS. RNA was extracted essentially as previously described (29) with some modifications as described below. TRI reagent (MRC) was added to cells at a ratio of 1 ml of TRI reagent to 1 × 10^7^ cells. Subsequently, for conventional RNA extraction, cell lysates in TRI reagent was extracted according to the manufacturer’s instruction. For improved RNA extraction, cell lysates in TRI reagent were diluted 1 × 10^6^ cells/ml using TRI reagent and passed 100 times through a 20-gauge needle. Subsequently, total RNA was extracted according to the manufacturer’s instructions. An aliquot of 1 μg of purified RNA was subjected to ribosomal RNA depletion using a Ribo-Zero Gold kit (Epicentre). Sequencing libraries were constructed from 100 ng total RNA using a Truseq stranded mRNA Library Prep kit (Illumina) without the poly(A) selection step. Subsequently, sequencing was performed using a Hiseq3000 (Illumina) with the 36-bp single-end or 101-bp paired-end method.

### RNA-seq analysis

Paired-end reads were trimmed using cutadapt (v3.5)(30) with the following parameters: -a AGATCGGAAGAGCACACGTCTGAACTCCAGTCAC -A AGATCGGAAGAGCGTCGTGTAGGGAAAGAGTGTA --overlap 5 --trim-n --max-n 1 --minimum-length 50:50. For single-end reads, the adapter-removal step was skipped. The human genome sequence (hg38) and gene annotation were downloaded from the GENCODE (v43) project (31). Abundant RNAs were extracted from the gene annotation according to the criteria of transcript length less than 200 nt or transcript biotype containing tRNA, miRNA, snoRNA, rRNA. First, the reads were mapped to the abundant RNAs using STAR (v2.7.10a) (32) when multi-mapped reads were allowed. Thereafter, the unmapped reads were mapped to the genome using STAR with the following parameter: --outFilterMultimapNmax 1. Transcript/exon/intron FPKM (fragments per kilobase of exon per million mapped reads) was estimated using the StringTie (v2.2.1) (33) quantification mode (-e) with default parameters.

The reference transcriptome for quantification (FPKM and read count) was constructed in the following steps. For each gene, all annotated exons were collapsed and merged to a unique set of exons that constitutes a single transcript (hereinafter referred to as “representative transcript”). RNA-seq, obtained using the improved RNA extraction method, was used to assess intron retention and to assemble novel transcripts from intergenic regions. To assess the intron retention for a representative transcript, the retention score (RS) for the i-th intron was calculated as follows:

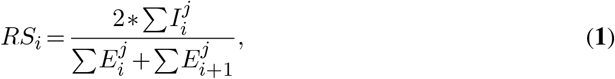

where *E* and *I* indicate the FPKM of exon and intron, respectively. *j∈*{A10, A549, HAP1, HEK, HeLa}. We defined a retained intron as its FPKM *>* 0.1 and RS *>* 0.1. All retained introns in a representative transcript were merged with it to form another one that we refer to as an “intron-retaining transcript”. Note that a gene may hold both a representative transcript and an intron-retaining transcript in the reference transcriptome. For each sample, StringTie assembles transcripts based on mapped reads with the following parameters: --rf. The transcripts obtained from all samples were grouped by forward and reverse strands and then merged separately using StringTie merge mode (--merge) with default parameters. Finally, the two groups of transcripts were concatenated into the reference transcriptome. After removal of those transcripts that overlap with the representative transcripts and their 5000 nt upstream and downstream regions, we obtained “intergenic transcripts”. The above mentioned representative transcripts, intron-retaining transcripts, and intergenic transcripts were combined into the complete reference transcriptome.

For the differential expression analysis between conditions (the conventional RNA extraction and the improved extraction), the transcript-level count of mapped reads was estimated using featureCounts (v2.0.1) (34) with the following parameters: -O -s 2. Then, the fold change (FC) and the p-value of transcript was calculated using edgeR (v3.28.1) (35). To apply edgeR analysis on the partial datasets without biological replicates, we derived an average dispersion of 0.125 from the other datasets as an empirical parameter. For each cell line, transcripts were categorized based on the following criteria: those with average FPKM *<* 0.5 were labeled as low expressed (Low); those with average FPKM *≥* 0.5, p-value *<* 0.05, and log_2_FC *>* 0 were categorized as up-regulated (Up); those with average FPKM *≥* 0.5, p-value *<* 0.05, and log_2_FC *<* 0 were categorized as down-regulated (Down); and the remaining transcripts were considered non-significant (NS). We consider that a transcript categorized as “Up” in a cell line indicates that it exhibits semi-extractability for the cell line. Transcripts can be labeled “common”, “specific”, or “switch” based on their categorization in A10, A549, HAP1, HEK, and HeLa cell lines. If a transcript is categorized as one category in all five cell lines, it is labeled “common” as either “Up common”, “Down common”, or “Low common”. If a transcript is categorized as two categories in all five cell lines, with one being a Low category, it is labeled “specific”. Transcripts that do not fall into the above categories are labeled as “switch”, for example, “Up-NS switch” or “Down-NS switch”. The label “Up common” indicates that the transcript is consistently semi-extractable in all five cell lines, while “Up specific” means it is specifically expressed in some cell lines and is semi-extractable in all of them. For subsequent meta-analysis, we defined transcripts labeled as “Up common” or “Up specific” as a set of semi-extractable RNAs (denoted as “SE”), and transcripts labeled as “NS common” as a set of extractable RNAs (denoted as “EX”). Note that in SE or EX, the representative transcript is removed when both the representative transcript and the intron-retaining transcript of a gene are present. Additionally, the reference transcriptome (including representative transcripts, intron-retaining transcripts, and intergenic transcripts) were prepared as background controls (denoted as “BG”).

The UCSC genome browser (36) was used to visualize the reference transcriptome and read coverage. To visualize read coverage, the mapped reads in BAM format were indexed using Samtools (v1.14) (37) and then converted to bigWig format with bamCoverage (v3.5.1) (38) using the following parameters: --filterRNAstrand forward/reverse --scaleFactor 1/-1 -bs 1 --normalizeUsing RPKM.

### Chromatin state

The chromatin states of HeLa cells were downloaded from the ENCODE (39) project (http://hgdownload.cse.ucsc.edu/ goldenpath/hg19/encodeDCC/wgEncodeAwgSegmentation/ wgEncodeAwgSegmentationChromhmmHelas3.bed.gz). These chromatin states were predicted using a trained ChromHMM (40) model based on multiple chromatin datasets, including ChIP-seq data for various histone modifications. The annotations of chromatin states that were on hg19 were remapped to hg38 using the pyliftover package (https://github.com/konstantint/pyliftover). The chromatin state prefixes were re-annotated as follows: (i) Active Promoter: Tss and TssF; (ii) Promoter Flanking: PromF; (iii) Inactive Promoter: PromP; (iv) Candidate Strong enhancer: Enh and EnhF; (v) Candidate Weak enhancer/DNase: EnhWF, EnhW, DNaseU, DNaseD; (vi) Distal CTCF/Candidate Insulator: CtrcfO and Ctcf; (vii) Transcription associated: Gen5^*′*^, Elon, ElonW, Gen3^*′*^, Pol2, H4K20. (viii) Low activity proximal to active states: Low. (ix) Polycomb repressed: ReprD, Repr, and ReprW; and (x) Heterochromatin/Repetitive/Copy Number Variation: Quies, Art. The chromatin states were then intersected with semi-extractable and extractable RNAs using the BEDTools (41) intersect command.

### Subcellular localization

APEX-seq data for HEK cells were obtained from GSE116008. APEX-seq is an RNA sequencing method coupled with direct RNA proximity labeling (42). For each cell compartment, we measured the enrichment of a transcript in that compartment (termed subcellular localization) by calculating the fold-change in the abundance of that transcript between labeled and unlabeled libraries. Accordingly, the RNA-seq reads were first subjected to adapter trimming using Trimmomatic (v0.39) (43) with the following parameters: ILLUMINACLIP:adapter.fa:2:30:4 TRAILING:20 MINLEN:36. Then the reads were uniquely mapped to the human genome using STAR, and the transcript abundance was estimated using StringTie. Finally, subcellular localization (log2 fold-change in transcript abundance) was calculated using an in-house script. For a transcript, a higher value of subcellular localization value indicates a higher enrichment in the corresponding cell compartment.

### Minimum free energy analysis

Using a transcript, subsequences of 300 nt length were extracted from its 5^*′*^ and 3^*′*^ ends. Transcripts less than 600 nt in length were removed beforehand. These subsequences were subjected to minimum free energy (MFE) calculations using RNAfold (v2.5.0) (44) with default parameters. Generally, a lower MFE value indicates a more stable RNA structure.

### RNA-chromatin interaction

*In situ* mapping of RNA-Genome Interactome (iMARGI) data of HEK cells were downloaded from GSM3478205. iMARGI is a DNA sequencing method based on RNA-DNA proximity ligation *in situ* inside an intact nucleus (45). The genomic coordinates of the RNA ends in the RNA-DNA interactions were extracted from the processed iMARGI data. The RNA ends were then intersected with transcripts using the BEDTools intersect command. To measure the extent to which a transcript interacts with chromatin, the fraction of transcript regions covered by iMARGI RNA ends was calculated. This fraction ranged from 0 to 1, with a higher fraction suggesting a more frequent interaction between the transcript and chromatin.

### RNA-RNA interaction

RNA interaction hubs (termed “hub RNAs”) were derived from RIC-seq (RNA *in situ* conformation sequencing) (46) and PARIS (psoralen analysis of RNA interactions and structures) (47). Hub RNAs exhibited stronger trans-interactions than other RNAs. RIC-seq can detect protein-mediated RNA-RNA interactions, while PARIS can directly identify RNA duplexes in trans across the transcriptome. The hub RNAs defined by RIC-seq were obtained from the previous study (46). Hub RNAs defined by PARIS were downloaded from the RISE database (48) and reserved for RNAs associated with more than twenty RNAs simultaneously in HeLa cells. Only 9,712 expressed genes (average FPKM *≥* 0.5, removal of intergenic genes and genes without gene names) in HeLa cells considered when computing the enrichment of overlap.

### Repeat density

The genomic coordinates of the repeat sequences were extracted using RepeatMasker (hg38, repeat library 20140131; https://www.repeatmasker.org/species/hg.html). For a transcript, BEDtools was used for intersection with repeat sequences. The fraction of the transcript that overlapped with repeat sequences, termed repeat density, was then calculated using an in-house script.

### Sequence motif analysis

Human RBP-binding sequence motifs (position weight matrix format) were downloaded from the CISBP-RNA database (http://cisbp-rna.ccbr.utoronto.ca; accessed on March 12, 2022). For a transcript, FIMO (v5.4.1) (49) scanned RBP-binding sites based on the above motifs using the following parameters: --norc --thresh 0.01 --motif-pseudo 0.1 --max-stored-scores 100000000. Given a transcript, the binding score of a certain RBP was defined as the number of binding sites of this RBP normalized by the transcript length. The binding preference of an RBP for semi-extractable RNAs is the ratio (log2 scale) of its average binding score to that of extractable RNAs.

### K-mer analysis

Semi-extractable RNAs were functionally classified using the k-mer content-based SEEKR (50) algorithm. First, seekr kmer counts was used to count the frequency of k-mer occurrence with the following parameter: -k 6. Thereafter, seekr pearson was used to calculate the similarity matrix. Finally, seekr graph segmented the RNA sequences into different communities based on the similarity matrix described above and the following parameters: 0.19 --louvain. The network graph of the semi-extractable RNAs was visualized using Gephi (v0.9) (51) with a Yifan Hu proportional layout.

### Gene ontology analysis

Gene ontology (GO) analysis of the semi-extractable genes across the five cell lines was conducted using g:Profiler (version: e105 eg52 p16 e84549f) (52). Statistical domain scope: only annotated genes; significance threshold: Bonferroni correction; user threshold: 0.001.

### Data availability

The conventional and semi-extractable RNA-seq of A10, A549, HAP1, and HEK cells have been deposited in the DDBJ Sequence Read Archive (DRA, https://www.ddbj.nig.ac.jp) under accession numbers DRA009793, DRA012807, DRA012808, DRA012810, and DRA014991. Published RNA-seq of HeLa cells were retrieved from Gene Expression Omnibus (GEO, https://www.ncbi.nlm.nih.gov/geo) under accession number GSE80589.

## RESULTS

### Reference transcriptome for identifying semi-extractable RNAs

To build a reference transcriptome for identification of the semi-extractable RNAs and intron retention, transcriptome reconstruction was performed based on the RNA-seq data produced by the improved RNA extraction (Figure 1A). The rationale for this approach is based on our observation that numerous semi-extractable RNAs are not properly annotated in the existing public databases. For example, hundreds of readthrough downstream-of-gene (DoG) transcripts were discovered to be semi-extractable and reported in another study (53). Semi-extractable RNAs may be the products and intermediates of various steps (e.g., transcription, processing, and degradation) and thus contain intronic sequences or partially missing exonic sequences. Further, a semi-extractable RNA may not be available in the existing gene annotations, because it can derived from intergenic regions.

**Figure 1.**
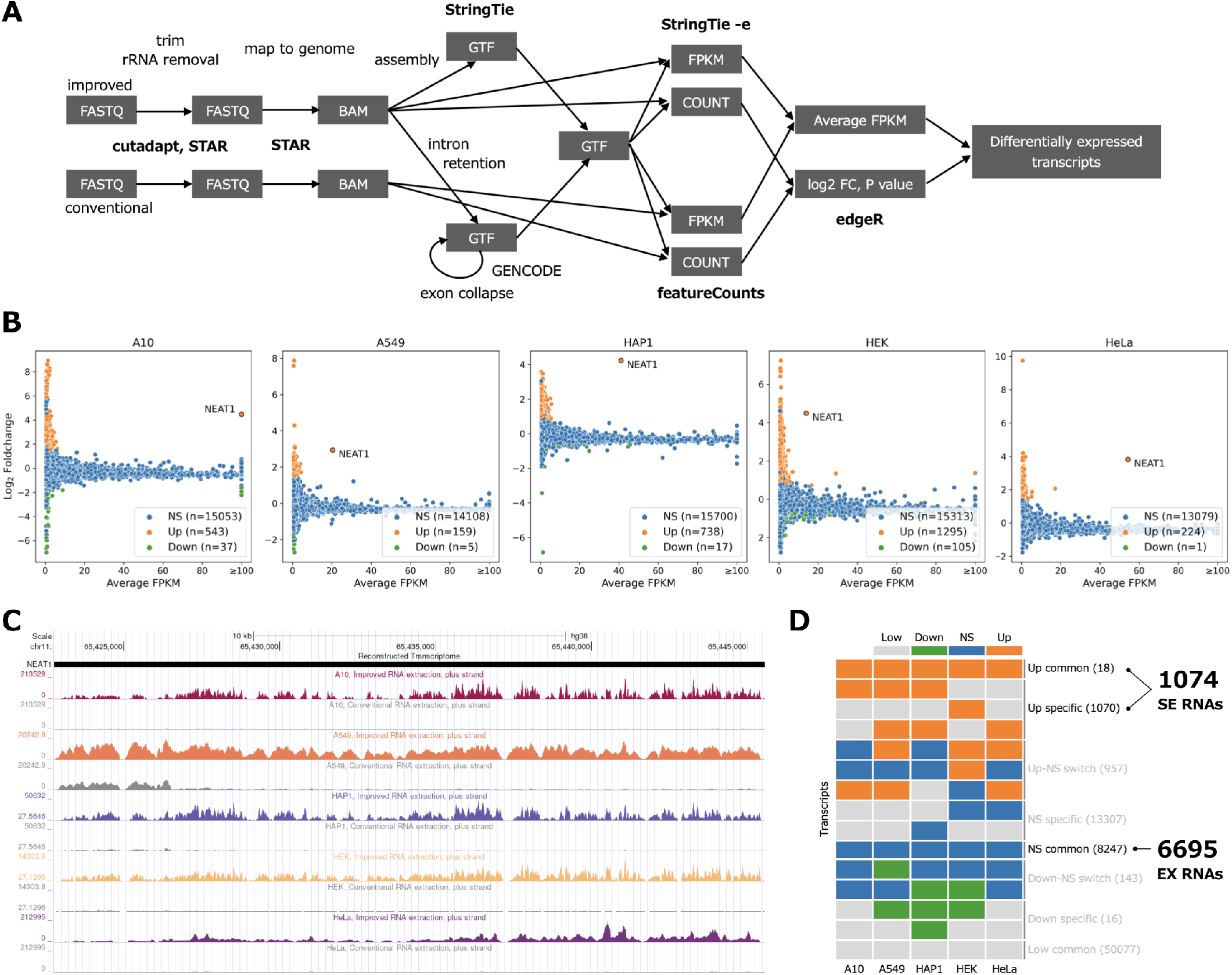
Identification of semi-extractable RNAs. **(A)** RNA-seq data analysis workflow. **(B)** Identification of semi-extractable/up-regulated RNAs (Up, orange), down-regulated RNAs (Down, green), and non-significant RNAs (NS, blue) from A10, A549, HAP1, HEK, and HeLa cells. **(C)** NEAT1 was simultaneously detected as a semi-extractable RNA in the five cells. **(D)** Schematic of defining semi-extractable RNAs across five cells. A total of 1,074 semi-extractable (SE) RNAs and 6,695 extractable (EX) RNAs were obtained after merging from the five cells. In the case that both representative transcript and intron-retaining transcript are present in a gene, the former is removed. Low: low-expressed, Down: down-regulated, NS: non-significant, Up: up-regulated. See “Materials and Methods” and Additional file 4 for details.

The multi-mapping reads were removed prior to evaluating intron retention. Most of the RNA-seq reads in this study were short single-ended (36 nt), maintaining an empirically low unique mapping rate (71% on average, see Additional file 1). Uniquely mapped reads (referred to as uniq-reads) have a higher confidence than multi-mapping reads of ambiguous origin. For single-ended reads, multi-mapping reads tended to cause higher read coverage over regions including simple repeat and/or low-complexity sequences (Additional file 2, bottom). Such ambiguous regions even confounded the surrounding high-confidence regions that were supported by uniq-reads. The above situation was not alleviated by applying longer paired-end reads (101 nt, Additional file 2, upper). Accordingly, we concluded that multi-mapping reads may lead to ambiguous transcriptome construction, especially in regions containing simple repeats and/or low-complexity sequences.

For each gene, all exons were collapsed to obtain a representative transcript. For the remaining intron(s), we applied the retention score (RS) to quantify its retention level based on the read coverage. The RS value reflects the expression ratio of the intron relative to its flanking exons. We combined the introns with RS values greater than 0.1 with the original representative transcript to obtain an intron-retaining transcript. In addition, for the intergenic region, we adopted a genome-based transcriptome assembly to construct intergenic transcripts. Finally, we obtained a reference transcriptome containing 57,001 representative transcripts, 13,702 intron-retaining transcripts and 3,132 intergenic transcripts (Additional file 3).

### A total of 1,074 semi-extractable RNAs were identified across five human cell lines

For each reference transcript, we quantified its semi-extractability using the expression increment obtained by the improved extraction method versus the conventional extraction method. A larger increment indicates higher semi-extractability of the transcript. We defined significant (p-value *<* 0.05) up-regulated transcripts as semi-extractable RNAs. Finally, 159–1,295 semi-extractable RNAs were identified from each of the five cell lines (Figure 1B, Additional file 5). NEAT1 lncRNA has been reported to be the most remarkable semi-extractable RNA in HeLa cells (29). This result was reproduced using HeLa cells, as shown in (Figure 1B-C). NEAT1 was found to exhibit consistent semi-extractability in four other cell lines. Moreover, the expression level of NEAT1 was almost the highest among all semi-extractable RNAs.

We proceeded to determine whether transcripts other than NEAT1 exhibited stable semi-extractability across various cell lines. We investigated the overlap between the semi-extractable RNAs identified in A10, A549, HAP1, HEK, and HeLa cells (Figure 1D). A total of 1,074 different semi-extractable (SE) RNAs were detected in the five cell lines. Of these genes, most were specifically expressed in only some of the five cell lines and exhibited semi-extractability. Interestingly, we discovered that eighteen transcripts, including ASAP1, DANT2, EXT1, FTX, IGF1R, LIMS1, NEAT1, PHF21A, PVT1, SCMH1, STRG.3024.1, TBL1X, TCF7L2, TVP23C-CDRT4, UBE2E2, ZCCHC7, ZFAND3, and ZSWIM6, exhibited consistent and stable semi-extractability in all cell lines (Figure 1D and Additional file 5). NEAT1, DANT2, FTX are long non-coding RNAs, STRG.3024.1 is a novel transcript identified in the intergenic region, and the remaining fourteen transcripts encode proteins. For control, we identified 6,695 extractable (EX) RNAs that were expressed in all five cell lines. These RNAs showed no significant difference in expression between the improved and conventional extraction methods (Figure 1D). In addition, all transcripts were prepared from the reference transcriptome as a background group (BG).

### Semi-extractable RNAs as a platform to provide RNA-RNA interactions

To investigate the distribution of semi-extractable RNAs in the chromatin, we compared their origins with the chromatin states downloaded from the ENCODE project (Table 1). Compared to extractable RNAs, semi-extractable RNAs were more enriched in repressed (polycomb repressed and low activity proximal to active states) and repetitive/heterochromatin (heterochromatin/repetitive/copy number variation) regions, with limited distribution in active promoter regions. In particular, there was a significant enrichment (SE/EX = 28.82) of semi-extractable RNAs in polycomb repressed regions. This enrichment could suggest that these genes are under regulatory control by Polycomb proteins. One possibility is that the semi-extractable genes are involved in processes that are not essential for cell survival. For example, genes that are only expressed during early development or in specialized cells might be more likely to be repressed by Polycomb proteins.

**Table 1.** Semi-extractable RNAs are preferentially transcribed from the repressed and heterochromatin/repetitive regions. The chromatin state in HeLa cells was annotated in advance using chromHMM and obtained from the ENCODE project. The percentages of various chromatin states in the transcribed regions of semi-extractable RNAs (%SE), extractable RNAs (%EX), and reference/background RNAs (%BG) were calculated separately. The ratio of %SE to %EX (SE/EX) and %BG (SE/BG) measures the transcriptional preference of semi-extractable RNAs in different chromatin states. Sorted by SE/EX column in descending order.

| ChromHMM states | %SE | %EX | %BG | SE/EX | SE/BG |
| --- | --- | --- | --- | --- | --- |
| Polycomb repressed | 4.85 | 0.20 | 4.41 | 23.82 | 1.10 |
| Heterochromatin/Repetitive/Copy Number Variation | 29.22 | 4.07 | 16.80 | 7.18 | 1.74 |
| Low activity proximal to active states | 50.44 | 31.02 | 44.67 | 1.63 | 1.13 |
| Candidate Weak enhancer/DNase | 3.74 | 3.56 | 4.91 | 1.05 | 0.76 |
| Distal CTCF/Candidate Insulator | 1.11 | 1.16 | 1.47 | 0.95 | 0.75 |
| Candidate Strong enhancer | 1.15 | 1.32 | 1.39 | 0.88 | 0.83 |
| Inactive Promoter | 0.07 | 0.18 | 0.26 | 0.41 | 0.28 |
| Promoter Flanking | 0.42 | 2.15 | 0.94 | 0.19 | 0.44 |
| Transcription associated | 7.99 | 46.37 | 18.92 | 0.17 | 0.42 |
| Active Promoter | 0.63 | 9.80 | 3.87 | 0.06 | 0.16 |

We proceeded to examine the subcellular localization of the semi-extractable RNAs. For each transcript, we calculated the degree of preference for nine subcellular fractions from publicly available APEX-seq data (42). Semi-extractable RNAs were significantly (p-value *<* 0.001) enriched in the nuclear compartments, including the nucleolus and nuclear lamina (Figure 2A). We further investigated the association of semi-extractable RNAs with chromatin using public iMARGI data (45). Semi-extractable RNAs were found to be disassociated from chromatin (Figure 2B). Overall, semi-extractable RNAs appear to be localized in the nucleus but isolated from the chromatin.

**Figure 2.**
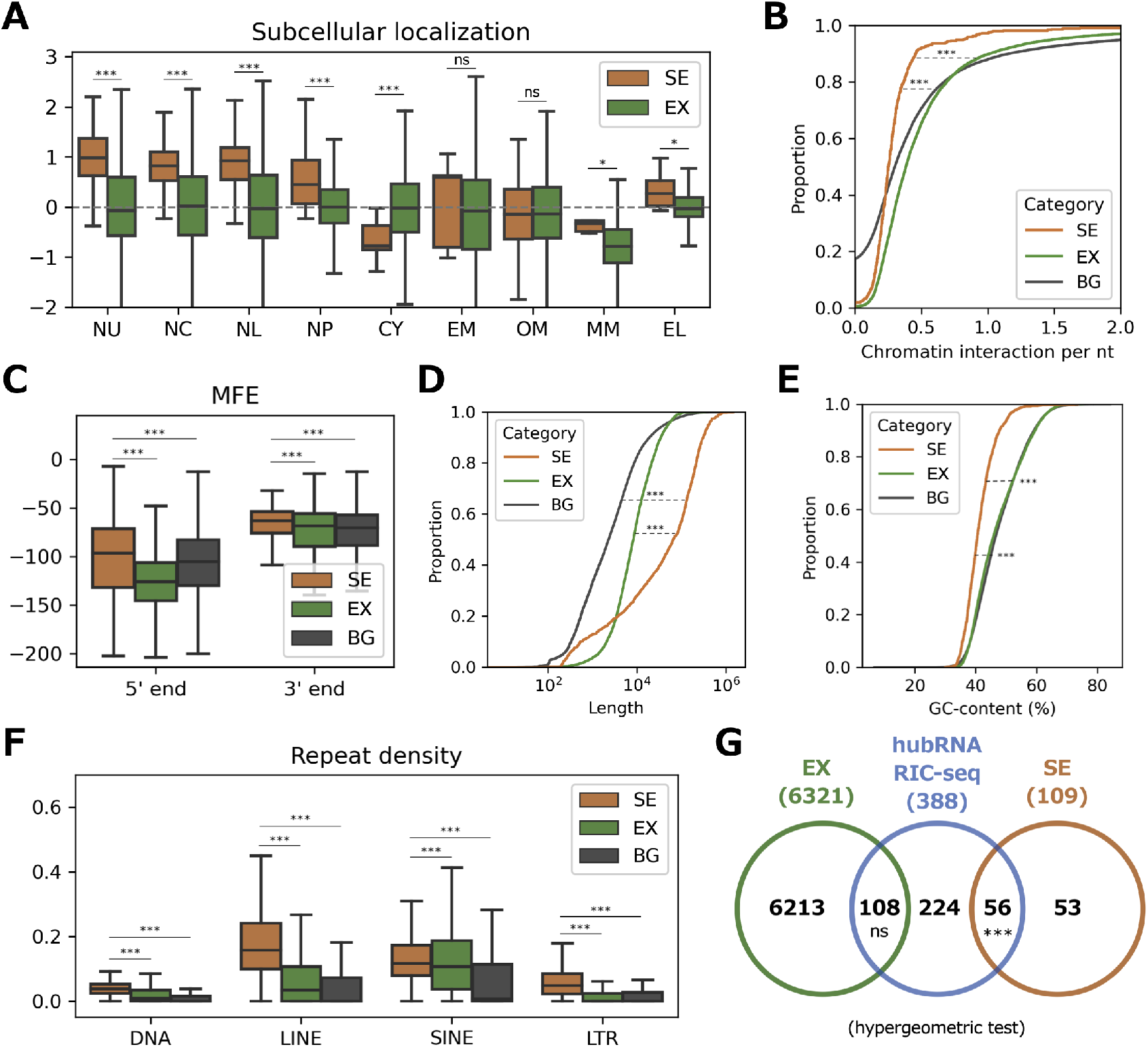
Characterization of semi-extractable RNAs. **(A)** Comparing subcellular RNA localization measured by APEX-seq fold changes in HEK cells. Increasing values indicate higher abundance in the corresponding subcellular fractions. NU: nucleolus, NC: nucleus, NL: nuclear lamina, NP: nuclear pore, CY: cytosol, EM: ER membrane, OM: outer mitochondrial membrane, MM: mitochondrial matrix, EL: ER lumen. **(B)** Chromatin-RNA interactions measured by iMARGI in HEK cells. **(C)** Comparing the minimum free energy (MFE) in the 5^*′*^ and 3^*′*^ end regions that are 300 nucleotides in length. MFE was calculated based on RNAfold. Cumulative density function analysis of **(D)** length in nucleotide, **(E)** G and C content. **(F)** Repeat elements are significantly enriched in semi-extractable RNAs. SINE: Short interspersed nuclear elements, DNA: DNA transposons, LINE: Long interspersed nuclear elements, LTR: Long terminal repeat. **(G)** Venn diagram analysis of semi-extractable RNAs and hub RNAs detected by RIC-seq in HeLa cells. ***: p-value *<* 0.001, **: p-value *<* 0.01,*: p-value *<* 0.05, ns: no significance (Wilcoxon rank-sum test is indicated if not otherwise specified). SE: semi-extractable RNAs, EX: extractable RNAs, BG: all background/reference RNAs.

NEAT1 forms paraspeckles through specific sequence features and an RNA-based interactome (26, 28). As NEAT1 was also identified as a consistent semi-extractable RNA aross cell lines in this study, we were curious whether semi-extractable RNAs possessed sequence characteristics similar to those of NEAT1. First, we determined whether the 5^*′*^ and 3^*′*^ ends of the semi-extractable RNAs had strong RNA structures to maintain RNA stability (28). We used the MFE of a sequence as a proxy for measuring the strength of the RNA structure. For an RNA sequence, a lower MFE indicates a higher propensity for strongly structured RNA. Surprisingly, the 5^*′*^ and 3^*′*^ ends of semi-extractable RNAs tended to have weak RNA structures (Figure 2C). In addition, we observed that the semi-extractable RNAs were significantly longer (Figure 2D) and with lower GC content (Figure 2E) than the extractable RNAs. Repeat elements (particularly LINEs) were significantly enriched in semi-extractable RNAs (Figure 2F). Based on the above observations, we hypothesized that semi-extractable RNAs are potential platforms for interactions with other RNAs. To test this hypothesis, we obtained 642 hub RNAs, which were detected to form protein-mediated RNA-RNA interactions with multiple RNAs from public RIC-seq data (46). Venn diagram analysis revealed that hub RNAs were significantly enriched (51.38%, 56 of 109) in the semi-extractable RNAs (Figure 2G). However, such enrichment of direct RNA-RNA interactions as defined in the PARIS public dataset were not observed (4.59%, 5 of 109, Additional file 6).

### Multifunctionality of the semi-extractable RNAs as reflected in clustered RBPs

We next explored the RBPs that bound to the semi-extractable RNAs. We downloaded the binding sequence motifs of 400 RBPs obtained by experimental validation from the CISBP-RNA database and used them to predict the binding preference of RBPs on semi-extractable RNAs. We found that RBPs that recognize AU-rich sequences were preferentially associated with semi-extractable RNAs (Figure 3, Additional file 7). AU-rich elements have been reported in the 3^*′*^ UTRs of many mRNAs and are associated with the regulation of RNA stability (54, 55). Interestingly, RBPs recognizing AU-rich elements were concentrated in the 5^*′*^ terminal regions of the semi-extractable RNAs (Figure 3), implying that AU-rich elements in semi-extractable RNAs may be involved in other uncovered functions.

**Figure 3.**
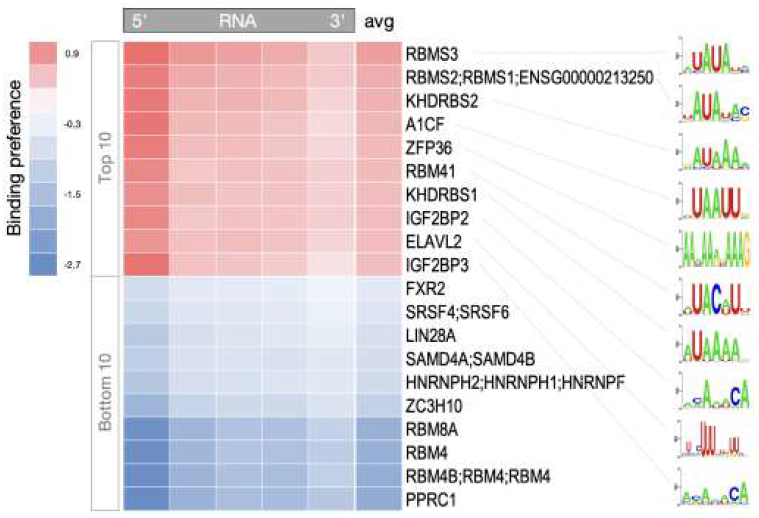
Motif enrichment analysis of semi-extractable RNAs. RBP binding preferences in different positional regions of semi-extractable RNAs, controlled with extractable RNAs. Here, x and y axes represent RNAs and RBPs, respectively. RNA was divided equally into five regions, and the RBP binding density within the regions was predicted using FIMO. The avg column indicates the average binding preference of RBP over the whole RNA. The result is sorted by the avg column. RBP binding sequence motifs are shown on the right. See Additional file 7 for details.

In addition, the reported paraspeckle RBPs enriched in NEAT1 (28) did not have a global binding preference for semi-extractable RNAs (Additional file 7). Hence, we hypothesized that the semi-extractable RNAs might contain functionally diverse RNAs, including a group that possesses functions similar to that of the NEAT1 constituent paraspeckles. Accordingly, we divided the semi-extractable RNAs into four potential functional groups/communities based on sequence similarity, determined by the Pearson”s correlation of k-mer profiles (Figure 4A). Among the eighteen stable semi-extractable RNAs, ten including NEAT1, PVT1 and others belonged to group 2 (Figure 4B). Furthermore, we examined the above four groups of semi-extractable RNAs for RBP-binding preference. Paraspeckle RBPs, such as FUS and NONO, preferentially bound to group 2 containing NEAT1 (Figure 4C). Theses results are consistent with those of a previous study (28).

**Figure 4.**
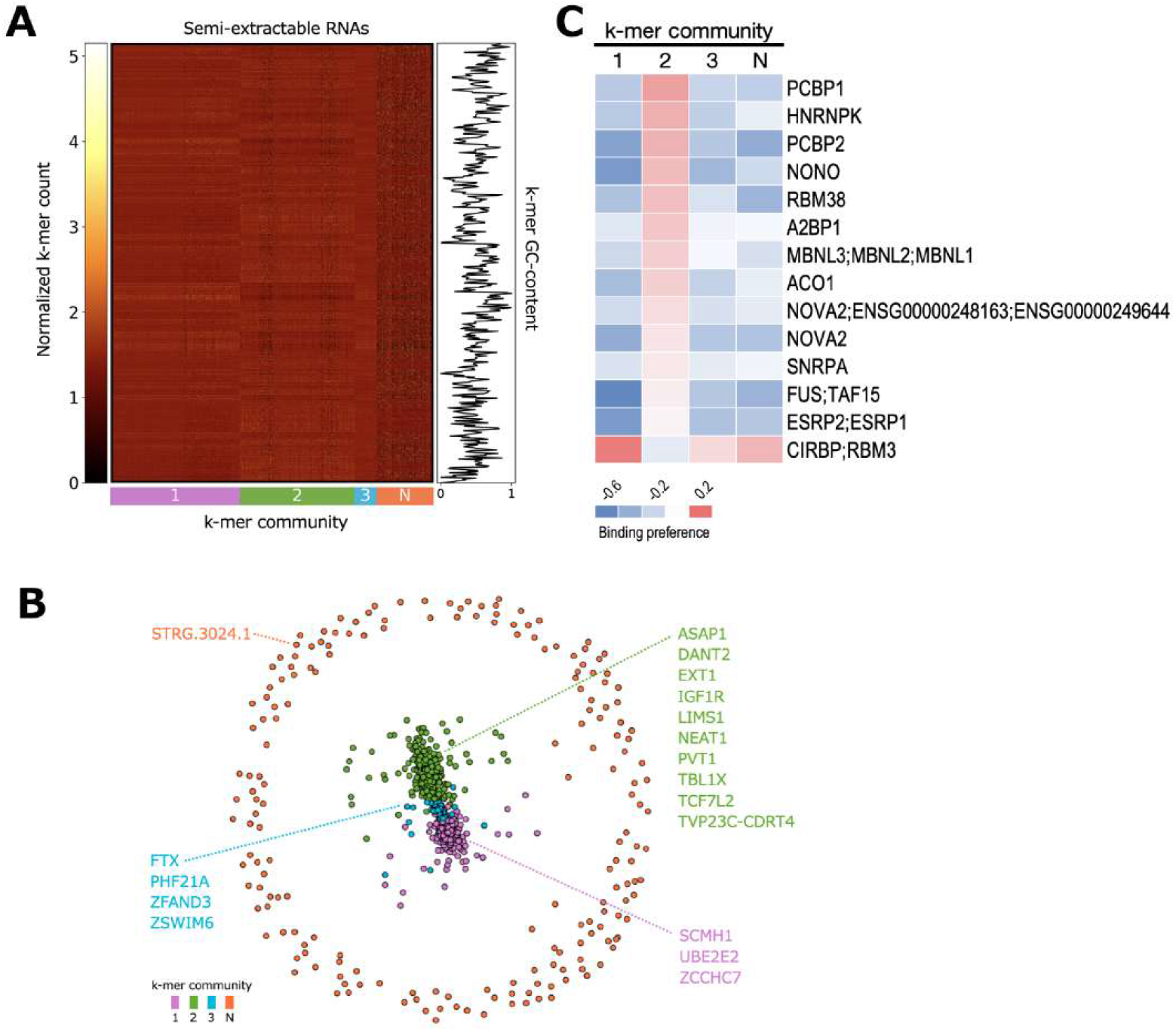
Clustering analysis of semi-extractable RNAs. **(A)** Louvain-assigned community of semi-extractable RNAs at k-mer length 6, with semi-extractable RNAs and k-mers on the x and y axes, respectively. Normalized k-mer count ranges from black (lowest) to yellow (highest). GC content of the k-mers is shown on the right panel. A side bar of the k-mer community is shown below the x axis. N means the null community. **(B)** Network graph of semi-extractable RNAs. RNA names are colored by their Louvain community assignment. Eighteen RNAs exhibiting consistent semi-extractability in five cell lines were labeled. **(C)** RBP-binding preference analysis was performed for each semi-extractable RNA community separately. Only the specific RBP binding preferences (positive and negative) for community 2 is shown. See Additional file 9 for details.

## DISCUSSION

In this study, 1,074 semi-extractable RNAs were systematically identified from five human cell lines, thereby providing an essential resource for studying RNA-centric phase separation. Biomolecular condensates without membranes are typically formed via phase separation in cells. Most previous studies have focused on the role of various proteins in forming phase-separated structures, and many proteins associated with phase separation have been explored (1, 13, 14, 15). However, numerous researchers have recently turned their attention to the role of RNA in phase separation (16, 17, 18, 19, 20, 21). NEAT1 has been reported to act as an architectural RNA to form a membrane-less condensate in the nucleus, called the paraspeckle (22, 23, 24, 25). Previous studies have experimentally verified that semi-extractable RNAs, including NEAT1, can induce the formation of nuclear bodies (29). Therefore, the RNAs contained in condensates could be poorly harvested by conventional RNA extraction and exhibited semi-extractability. Thus, the semi-extractable RNAs detected in this study may have been derived from various phase-separated condensates. Such semi-extractable RNAs may be segregated into condensates by specific biological functions. However, GO analysis showed that semi-extractable RNAs were involved in a broad range of biological processes (Additional file 8). We proposed the following two hypotheses to explain this result: First, the semi-extractable RNAs may be a mixture of RNAs derived from condensates with different biological functions. As semi-extractable RNAs can be further classified according to the type of condensates, the specific biological functions involving these RNAs could be identified. Second, semi-extractable RNAs may be involved in specific biological regulatory processes as RNA molecules, and these functions are not detectable by GO analysis based on protein function and phenotype annotation.

According to subcellular localization analysis of semi-extractable RNAs, semi-extractable RNAs were enriched in the nuclear compartments. This phenomenon is consistent with the previous observation that semi-extractable RNAs are primarily derived from the nuclear bodies (29); this may be due to the dynamic exchange of contents (3), including RNAs, among nucleolus, nucleus, nuclear lamina, and nuclear pore. The semi-extractable RNAs were divided into four groups that may perform different biological functions based on sequence similarity (Figure 4B). Among them, the semi-extractable RNAs in group 2, where NEAT1 is located, preferentially bind to some known paraspeckle RBPs (i.e., NONO, FUS) (Figure 4C), implying that this group of semi-extractable RNAs may possess similar functions to NEAT1 in constituting the granule backbone. PVT1, as a stable semi-extractable RNA in this group (Additional file 5), was observed to form a complex network as hub RNAs with other RNAs through RBP-mediated RNA-RNA interactions, which can form granule-like foci in the nucleus, and PVT1 foci do not intersect with known nuclear bodies (29, 46). PVT1 is a neighbor of the well-known oncogene, MYC, and has been reported to be involved in the regulation of cancer development (56, 57, 58, 59, 60, 61). The semi-extractable property of PVT1 implies a new aspect of phase separation for investigating its molecular regulation in the mechanism of tumorigenesis. FTX in group 3 is involved in X chromosome inactivation as a positive regulator of XIST (62). This function has been reported to depend on FTX transcription, rather than its RNA product (63). However, the semi-extractability of FTX suggests that its RNA product may be involved in X chromosome inactivation via intracellular condensates. XIST has been reported to form a phase-separated compartment by interacting with multiple RBPs (20, 64, 65). However, in this study, the XIST was not observed to be semi-extractable. ZCCHC7 in group 1 is involved in RNA quality regulation after translation into proteins (66), especially viral RNA degradation (67). The semi-extractability of ZCCHC7 implies that its RNA product may be harbored in the cell as biomolecular condensates, without being eagerly used for protein production, but can rapidly respond to the invasion of pathogenic RNAs.

Numerous repetitive sequences were identified in the semi-extractable RNAs (Figure 2F), which is not consistent with our speculations, as we discarded the multi-mapping reads that may result from repetitive sequences. There could be several potential reasons for these results. First, many reads may be mapped to nonrepetitive regions for repeat-containing RNAs, allowing the expression levels of these RNAs to be detected. Second, reads containing repetitive sequences may still be uniquely mapped owing to mutations or unique flanking sequences in the repeats. Third, the default parameters we use in the transcript assembly cause some of the read coverage areas that are close together to be merged, which likely facilitated inclusion of repetitive sequences. Consistently, many RNAs that contain repeats have been reported to be associated with phase separation. For example, CTN-RNA was found to be distributed in mouse paraspeckles. CTN-RNA contains three inverted repeats from SINE, which are thought to affect A-to-I editing and nuclear retention (68). CAG-repeat-containing RNA was observed to colocalize with nuclear speckles that sequester splicing factors under *in vitro* conditions (69). HSATIII lncRNAs mainly consist of primate-specific satellite III repeats, which form nuclear stress bodies under thermal stress conditions (70) and recruit specific proteins, such as heat shock factor, chromatin-remodeling complex, and splicing factors (21, 71, 72). The middle domain of NEAT1 contains repetitive sequences from LINE and SINE and this region recruits NONO dimers to trigger paraspeckle assembly (28). A systematic analysis of the potential role of repetitive sequences in the formation of RNA condensates could deepen our understanding of the biological mechanism of phase separation (73, 74).

Finally, we opted to discuss possible limitations and directions for further work in this study. First, due to the various biases of short read RNA-seq (e.g., RNA fragmentation, PCR amplification, and sequence context), the transcriptome may not be assembled accurately. We may consider adding nanopore direct RNA-seq data to assist in obtaining full-length reference transcripts (75). Second, we compared semi-extractable RNAs with public experimental data (e.g., iMARGI, RIC-seq, and APEX-seq). Of note, these experimental data involved conventional RNA extraction and thus may have lost the information on semi-extractable RNAs. Repeating the above experiments while applying improved RNA extraction is a necessary direction of work to be completed. Fourth, various stress conditions can induce the formation of different phase-separated condensates (76, 77, 78). Therefore, exploring RNA semi-extractability under various stress conditions is expected to provide important clues for our study of the potential function of phase separation in the cellular stress response (Additional files 5 and 10, under analysis). Subsequent efforts will focus on RNAs that exhibit semi-extractability under specific stimulus conditions. Finally, there is growing evidence that RNA post-transcriptional modifications can regulate the dynamics of phase separation (79, 80, 81). An interesting direction of research is to investigate whether RNA modifications are associated with the semi-extractability of RNAs.

## CONCLUSION

To the best of our knowledge, this study provides the first dataset of genome-wide semi-extractable RNAs across cell lines (Additional file 5). This resource is expected to guide the exploration of RNA-based phase separations. Future use of semi-extractable RNAs in conjunction with RNA-centric interactome (45, 46, 82, 83, 84) will shed light on the molecular basis of the RNA-induced phase separation within cells.

## ADDITIONAL FILES

- Additional file 1. RNA-seq mapping statistics.
- Additional file 2. Multi-mapping reads lead to ambiguous coverage.
- Additional file 3. Gene annotation file for the reference transcriptome used to identify semi-extractable RNAs.
- Additional file 4. Full list of semi-extractable (SE) RNAs and extractable (EX) RNAs.
- Additional file 5. Differential gene expression analysis across cell lines and stress conditions.
- Additional file 6. Venn diagram analysis of semi-extractable RNAs and hub RNAs detected by PARIS in HeLa cell.
- Additional file 7. RBP binding preferences of semi-extractable RNAs.
- Additional file 8. GO analysis for semi-extractable genes.
- Additional file 9. RBP binding preferences for semi-extractable RNA groups.
- Additional file 10. Semi-extractable RNAs under different stress conditions.

## ACKNOWLEDGEMENTS

Computations were partially performed on the NIG supercomputer at ROIS National Institute of Genetics.

## FUNDING

This work was supported by JST CREST [grant no. JPMJCR20E6], AMED [grant no. JP21ae0121049], and JSPS KAKENHI [grants nos. 20H00448, 21H05276, and 22K19293] to TH; AMED [grant nos. JP22ama121055, JP21ae0121049 and JP21gm0010008], JSPS KAKENHI [grant nos. 22H04925, 20H00624, 17K20032] to MH; JSPS KAKENHI [grants nos. 20K15784, 22K15093] to CZ.

### Conflict of interest statement

None declared.

---

**Additional file 1.**
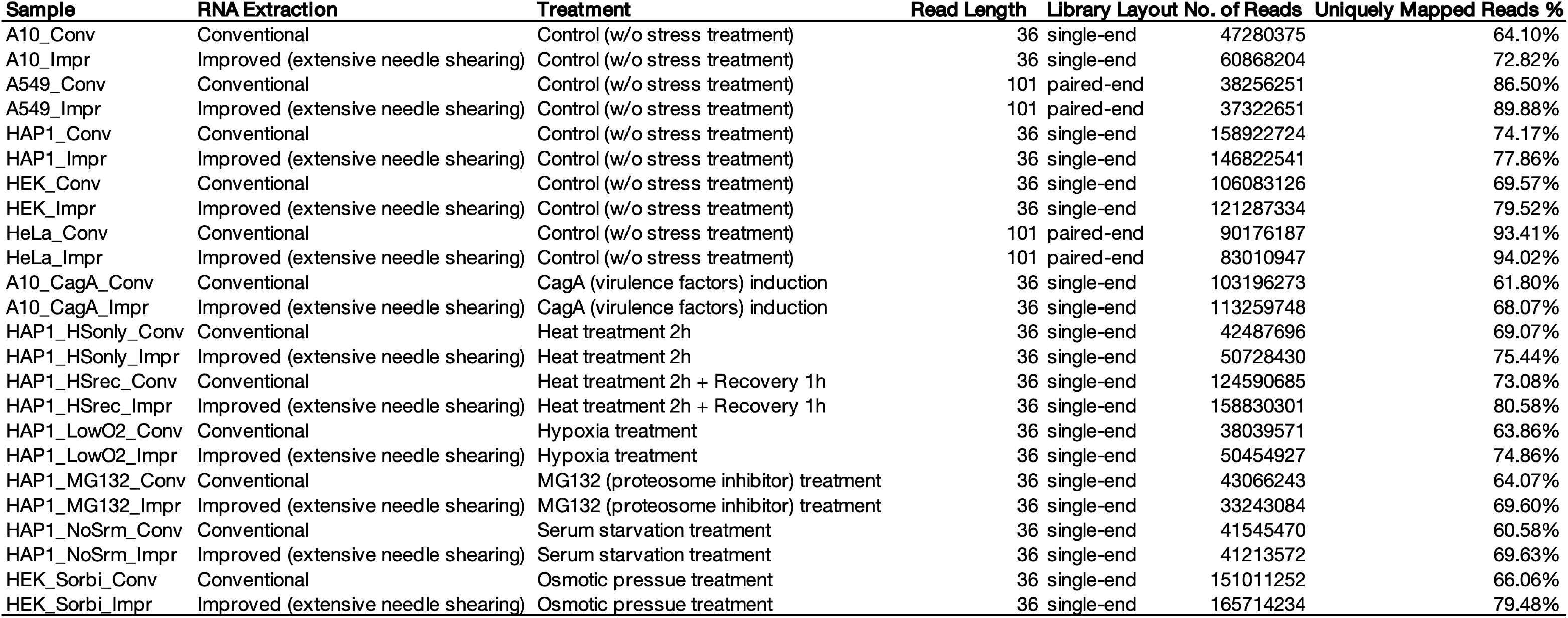
RNA-seq mapping statistics.

**Additional file 2.**
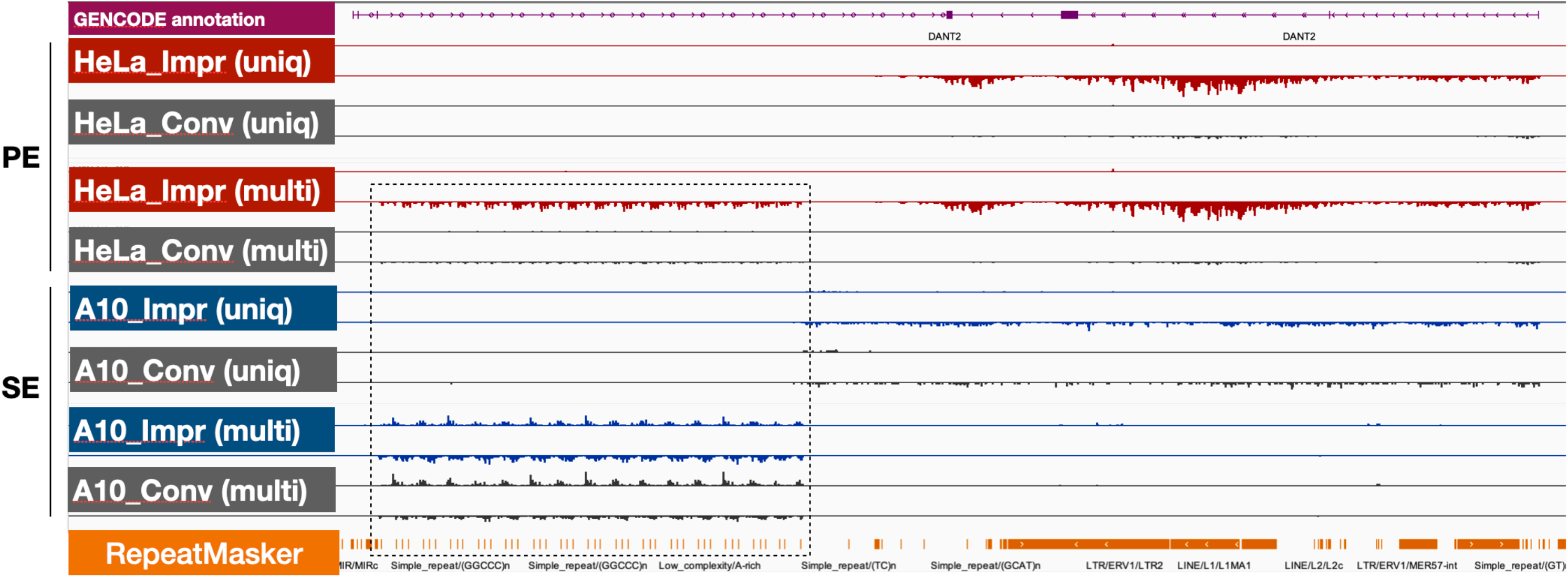
Multi-mapping reads lead to ambiguous coverage in intronic region. As an example of the DANT2 locus, either paired-end (PE, 101 nt) or single-end reads (SE, 36 nt), multiple mapped reads (multi) resulted in a read coverage of simple repeat/low complexity (dashed box) in intronic region. Read mapping were performed with STAR. Impr: improved RNA extraction, Conv: conventional RNA extraction.

**Additional file 3.**
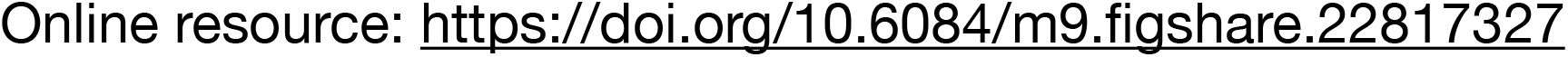
Gene annotation file for the reference transcriptome used to identify semi-extractable RNAs. A GTF (General Transfer Format) format file containing 7,001 representative transcripts, 13,702 intron-containing transcripts (transcript names with “_RI” suffix) and 3,132 intergenic transcripts (transcript names with “STRG.” prefix) obtained in this study.

**Additional file 4.**
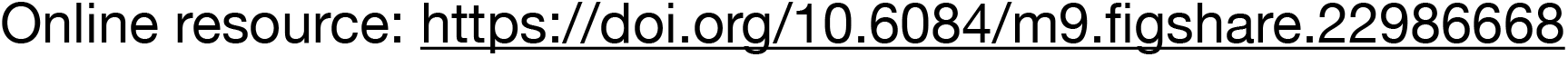
Full list of semi-extractable (SE) RNAs and extractable (EX) RNAs.

**Additional file 5.**
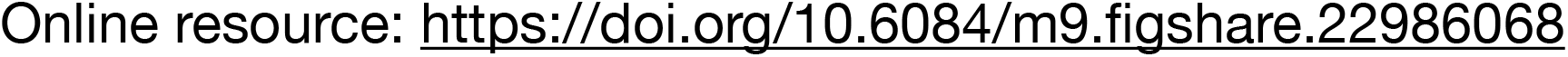
Differential gene expression analysis across cell lines and stress conditions.

**Additional file 6.**
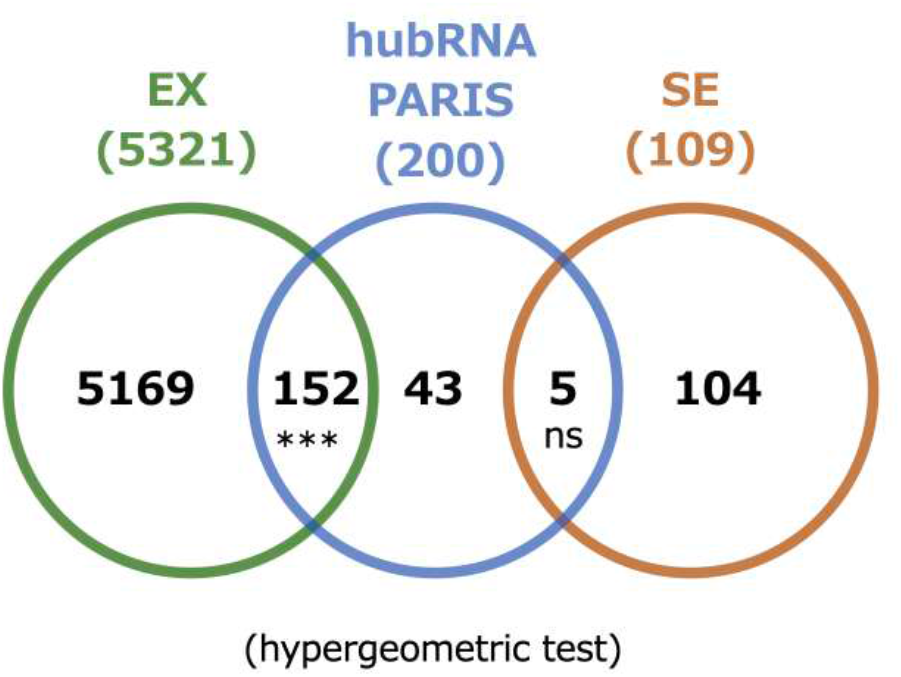
Venn diagram analysis of semi-extractable RNAs and hub RNAs detected by PARIS in HeLa cell. ***: p-value < 0.001, **: p-value < 0.01, *: p-value < 0.05, ns: no significance. SE: semi-extractable RNAs, EX: extractable RNAs.

**Additional file 7.**
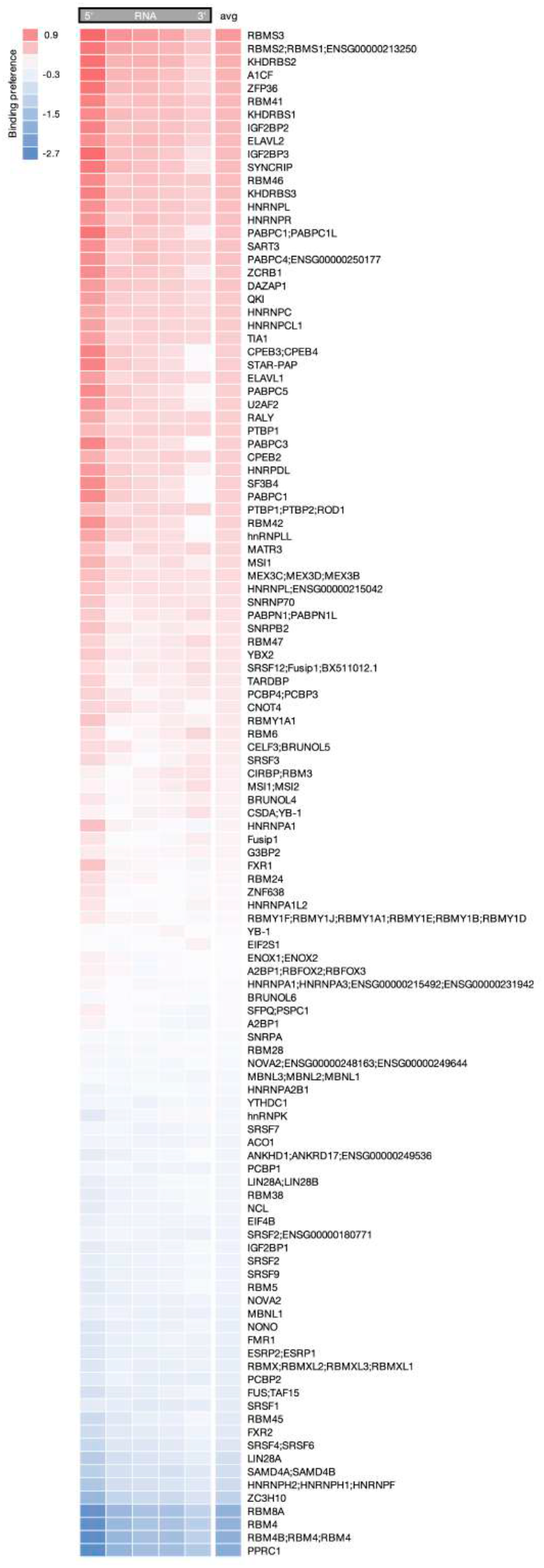
RBP binding preferences of semi-extractable RNAs.

**Additional file 8.**
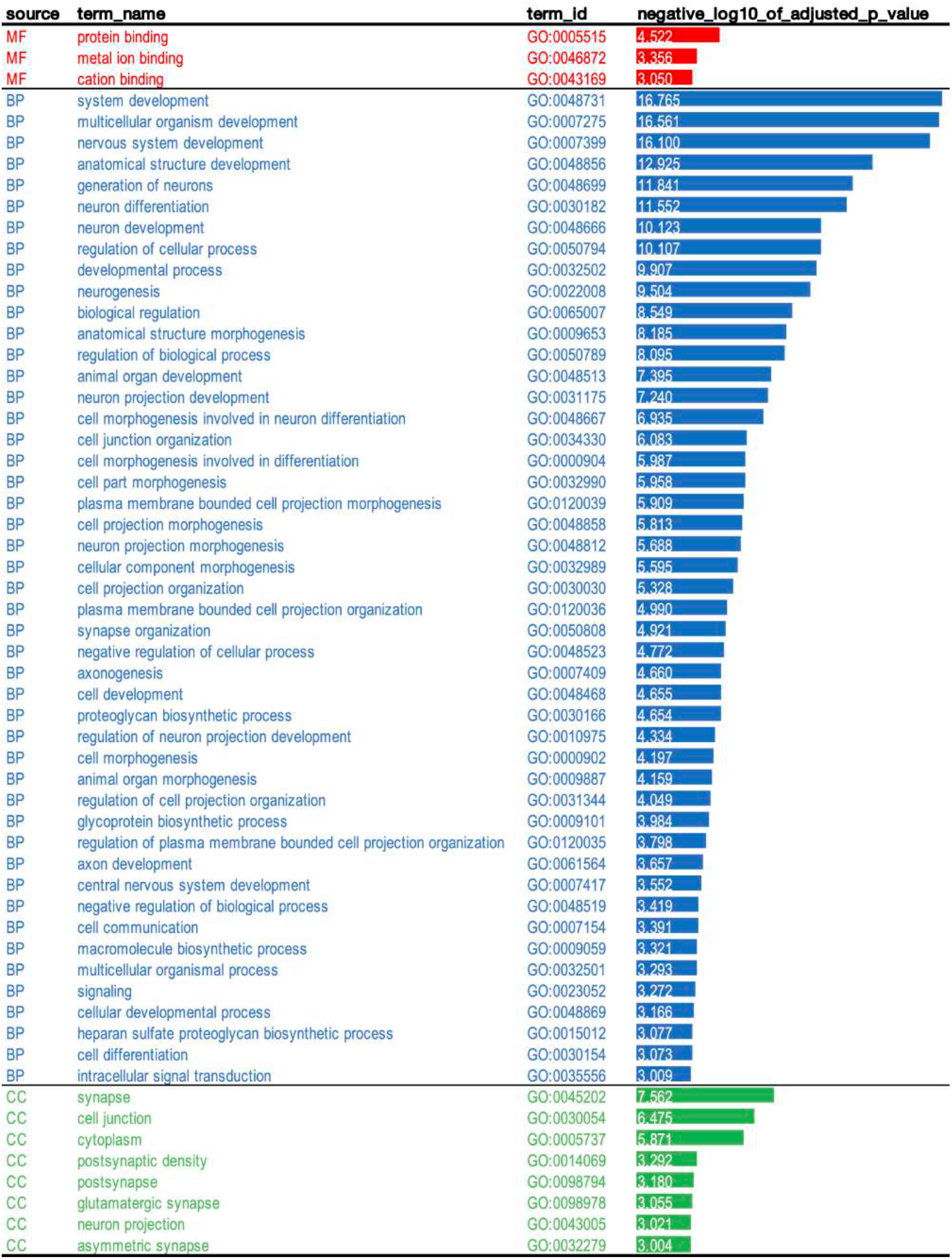
GO analysis for semi-extractable genes. GO analysis was conducted for 714 semi-extractable genes with g:Profiler. MF for molecular function, BP for biological process, and CC for cellular component

**Additional file 9.**
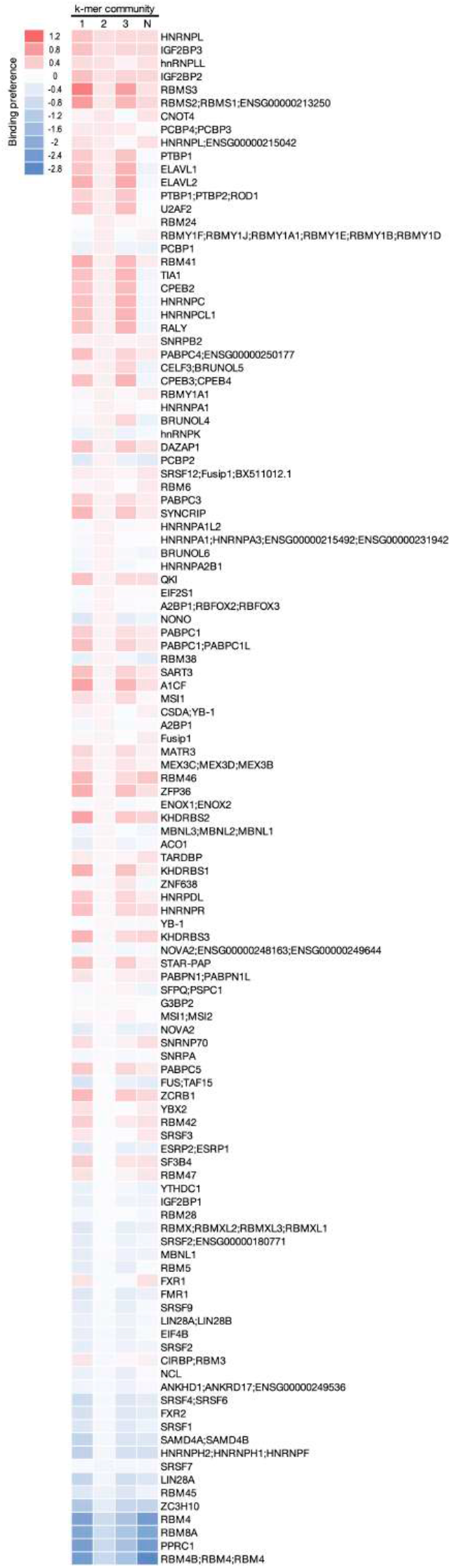
RBP binding preferences for semi-extractable RNA groups.

**Additional file 10.**
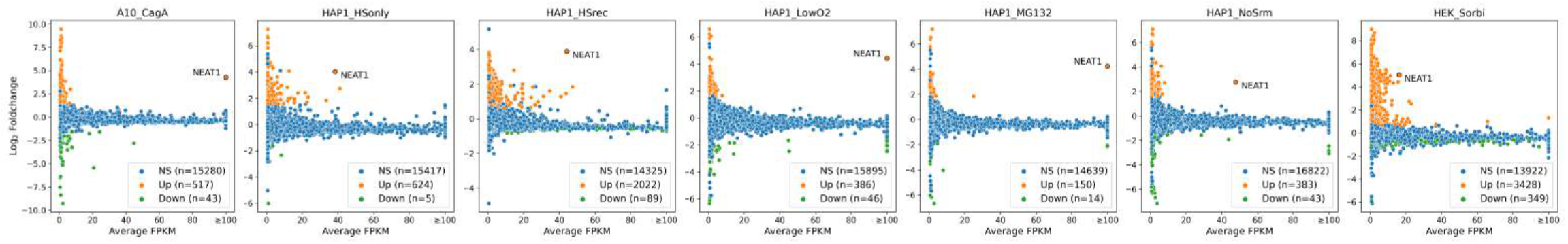
Semi-extractable RNAs under different stress conditions. Semi-extractable/up-regulated RNAs (Up, orange) and down-regulated RNAs (Down, green) were identified under seven stress conditions, respectively. CagA: CagA (virulence factors) induction. HSonly: heat treatment 2 hours. HSrec: heat treatment 2 hours and recovery 1 hour. LowO2: hypoxia treatment. MG132: MG132 (proteasome inhibitor) treatment. NoSrm: serum starvation treatment. Sorbi: osmotic pressure treatment.

## Notes

### Competing Interest Statement

The authors have declared no competing interest.

### Summary of Updates

Main text, tables, and figures updated; Additional files updated.

## REFERENCES

1. Salman F Banani, Hyun O Lee, Anthony A Hyman, and Michael K Rosen. Biomolecular condensates: organizers of cellular biochemistry. Nature Reviews Molecular Cell Biology, 18(5):285–298, 2017.

2. Gregory L Dignon, Robert B Best, and Jeetain Mittal. Biomolecular phase separation: From molecular driving forces to macroscopic properties. Annual Review of Physical Chemistry, 71:53, 2020.

3. Diana M Mitrea and Richard W Kriwacki. Phase separation in biology; functional organization of a higher order. Cell Communication and Signaling, 14(1):1–20, 2016.

4. Andrew S Lyon, William B Peeples, and Michael K Rosen. A framework for understanding the functions of biomolecular condensates across scales. Nature Reviews Molecular Cell Biology, 22(3):215–235, 2021.

5. Clifford P Brangwynne, Christian R Eckmann, David S Courson, Agata Rybarska, Carsten Hoege, Jöbin Gharakhani, Frank Jülicher, and Anthony A Hyman. Germline P granules are liquid droplets that localize by controlled dissolution/condensation. Science, 324(5935):1729–1732, 2009.

6. Mario Cioce and Angus I Lamond. Cajal bodies: a long history of discovery. Annual Review of Cell and Developmental Biology, 21:105, 2005.

7. Archa H Fox and Angus I Lamond. Paraspeckles. Cold Spring Harbor Perspectives in Biology, 2(7):a000687, 2010.

8. Angus I Lamond and David L Spector. Nuclear speckles: a model for nuclear organelles. Nature Reviews Molecular Cell Biology, 4(8):605–612, 2003.

9. David SW Protter and Roy Parker. Principles and properties of stress granules. Trends in Cell Biology, 26(9):668–679, 2016.

10. Bin Wang, Lei Zhang, Tong Dai, Ziran Qin, Huasong Lu, Long Zhang, and Fangfang Zhou. Liquid–liquid phase separation in human health and diseases. Signal Transduction and Targeted Therapy, 6(1):1–16, 2021.

11. Sohum Mehta and Jin Zhang. Liquid–liquid phase separation drives cellular function and dysfunction in cancer. Nature Reviews Cancer, 22(4):239–252, 2022.

12. Xuhui Tong, Rong Tang, Jin Xu, Wei Wang, Yingjun Zhao, Xianjun Yu, and Si Shi. Liquid–liquid phase separation in tumor biology. Signal Transduction and Targeted Therapy, 7(1):1–22, 2022.

13. Vladimir N Uversky. Intrinsically disordered proteins in overcrowded milieu: Membrane-less organelles, phase separation, and intrinsic disorder. Current Opinion in Structural Biology, 44:18–30, 2017.

14. Anastasia C Murthy, Gregory L Dignon, Yelena Kan, Gül H Zerze, Sapun H Parekh, Jeetain Mittal, and Nicolas L Fawzi. Molecular interactions underlying liquid-liquid phase separation of the FUS lowcomplexity domain. Nature Structural & Molecular Biology, 26(7):637–648, 2019.

15. Qian Li, Xi Wang, Zhihui Dou, Weishan Yang, Beifang Huang, Jizhong Lou, and Zhuqing Zhang. Protein databases related to liquid–liquid phase separation. International Journal of Molecular Sciences, 21(18):6796, 2020.

16. Man Wu, Guang Xu, Chong Han, Peng-Fei Luan, Yu-Hang Xing, Fang Nan, Liang-Zhong Yang, Youkui Huang, Zheng-Hu Yang, Lin Shan, et al. lncRNA SLERT controls phase separation of FC/DFCs to facilitate Pol I transcription. Science, 373(6554):547–555, 2021.

17. Rena Onoguchi-Mizutani, Yoshitaka Kirikae, Yoko Ogura, Tony Gutschner, Sven Diederichs, and Nobuyoshi Akimitsu. Identification of a heat-inducible novel nuclear body containing the long noncoding RNA MALAT1. Journal of Cell Science, 134(10):jcs253559, 2021.

18. Marina Garcia-Jove Navarro, Shunnichi Kashida, Racha Chouaib, Sylvie Souquere, Gérard Pierron, Dominique Weil, and Zoher Gueroui. RNA is a critical element for the sizing and the composition of phase-separated RNA–protein condensates. Nature Communications, 10(1):1–13, 2019.

19. Tetsuro Hirose, Tomohiro Yamazaki, and Shinichi Nakagawa. Molecular anatomy of the architectural NEAT1 noncoding RNA: The domains, interactors, and biogenesis pathway required to build phase-separated nuclear paraspeckles. Wiley Interdisciplinary Reviews: RNA, 10(6):e1545, 2019.

20. Amy Pandya-Jones, Yolanda Markaki, Jacques Serizay, Tsotne Chitiashvili, Walter R Mancia Leon, Andrey Damianov, Constantinos Chronis, Bernadett Papp, Chun-Kan Chen, Robin McKee, Xiao-Jun Wang, Anthony Chau, Shan Sabri, Heinrich Leonhardt, Sika Zheng, Mitchell Guttman, Douglas L Black, and Kathrin Plath. A protein assembly mediates Xist localization and gene silencing. Nature, 587(7832):145–151, 2020.

21. Kensuke Ninomiya, Shungo Adachi, Tohru Natsume, Junichi Iwakiri, Goro Terai, Kiyoshi Asai, and Tetsuro Hirose. LncRNA-dependent nuclear stress bodies promote intron retention through SR protein phosphorylation. The EMBO Journal, 39(3):e102729, 2020.

22. Takeshi Chujo, Tomohiro Yamazaki, and Tetsuro Hirose. Architectural RNAs (arcRNAs): A class of long noncoding rnas that function as the scaffold of nuclear bodies. Biochimica et Biophysica Acta (BBA)-Gene Regulatory Mechanisms, 1859(1):139–146, 2016.

23. Yasnory TF Sasaki, Takashi Ideue, Miho Sano, Toutai Mituyama, and Tetsuro Hirose. Men ε/β noncoding RNAs are essential for structural integrity of nuclear paraspeckles. Proceedings of the National Academy of Sciences, 106(8):2525–2530, 2009.

24. Christine M Clemson, John N Hutchinson, Sergio A Sara, Alexander W Ensminger, Archa H Fox, Andrew Chess, and Jeanne B Lawrence. An architectural role for a nuclear noncoding RNA: NEAT1 RNA is essential for the structure of paraspeckles. Molecular Cell, 33(6):717–726, 2009.

25. Hongjae Sunwoo, Marcel E Dinger, Jeremy E Wilusz, Paulo P Amaral, John S Mattick, and David L Spector. Men ε/β nuclear-retained non-coding RNAs are up-regulated upon muscle differentiation and are essential components of paraspeckles. Genome Research, 19(3):347–359, 2009.

26. Audrey Jacq, Denis Becquet, Séverine Guillen, Bénédicte Boyer, Maria-Montserrat Bello-Goutierrez, Jean-Louis Franc, and Anne-Marie François-Bellan. Direct RNA–RNA interaction between Neat1 and RNA targets, as a mechanism for RNAs paraspeckle retention. RNA Biology, 18(11):2016–2027, 2021.

27. Jason A West, Mari Mito, Satoshi Kurosaka, Toru Takumi, Chiharu Tanegashima, Takeshi Chujo, Kaori Yanaka, Robert E Kingston, Tetsuro Hirose, Charles Bond, Archa Fox, and Shinichi Nakagawa. Structural, super-resolution microscopy analysis of paraspeckle nuclear body organization. Journal of Cell Biology, 214(7):817–830, 2016.

28. Tomohiro Yamazaki, Sylvie Souquere, Takeshi Chujo, Simon Kobelke, Yee Seng Chong, Archa H Fox, Charles S Bond, Shinichi Nakagawa, Gerard Pierron, and Tetsuro Hirose. Functional domains of NEAT1 architectural lncRNA induce paraspeckle assembly through phase separation. Molecular Cell, 70(6):1038–1053, 2018.

29. Takeshi Chujo, Tomohiro Yamazaki, Tetsuya Kawaguchi, Satoshi Kurosaka, Toru Takumi, Shinichi Nakagawa, and Tetsuro Hirose. Unusual semi-extractability as a hallmark of nuclear body-associated architectural noncoding rna s. The EMBO Journal, 36(10):1447–1462, 2017.

30. Marcel Martin. Cutadapt removes adapter sequences from highthroughput sequencing reads. EMBnet. journal, 17(1):10–12, 2011.

31. Adam Frankish, Mark Diekhans, Irwin Jungreis, Julien Lagarde, Jane E Loveland, Jonathan M Mudge, Cristina Sisu, James C Wright, Joel Armstrong, If Barnes, Andrew Berry, Alexandra Bignell, Carles Boix, Silvia Carbonell Sala, Fiona Cunningham, Tomás Di Domenico, Sarah Donaldson, Ian T Fiddes, Carlos García Girón, Jose Manuel Gonzalez, Tiago Grego, Matthew Hardy, Thibaut Hourlier, Kevin L Howe, Toby Hunt, Osagie G Izuogu, Rory Johnson, Fergal J Martin, Laura Martínez, Shamika Mohanan, Paul Muir, Fabio C P Navarro, Anne Parker, Baikang Pei, Fernando Pozo, Ferriol Calvet Riera, Magali Ruffier, Bianca M Schmitt, Eloise Stapleton, Marie-Marthe Suner, Irina Sycheva, Barbara Uszczynska-Ratajczak, Maxim Y Wolf, Jinuri Xu, Yucheng T Yang, Andrew Yates, Daniel Zerbino, Yan Zhang, Jyoti S Choudhary, Mark Gerstein, Roderic Guigó, Tim J P Hubbard, Manolis Kellis, Benedict Paten, Michael L Tress, and Paul Flicek. Gencode 2021. Nucleic Acids Research, 49(D1):D916–D923, 2021.

32. Alexander Dobin, Carrie A Davis, Felix Schlesinger, Jorg Drenkow, Chris Zaleski, Sonali Jha, Philippe Batut, Mark Chaisson, and Thomas R Gingeras. STAR: ultrafast universal RNA-seq aligner. Bioinformatics, 29(1):15–21, 2013.

33. Sam Kovaka, Aleksey V Zimin, Geo M Pertea, Roham Razaghi, Steven L Salzberg, and Mihaela Pertea. Transcriptome assembly from long-read RNA-seq alignments with StringTie2. Genome Biology, 20(1):1–13, 2019.

34. Yang Liao, Gordon K Smyth, and Wei Shi. featurecounts: an efficient general purpose program for assigning sequence reads to genomic features. Bioinformatics, 30(7):923–930, 2014.

35. Mark D Robinson, Davis J McCarthy, and Gordon K Smyth. edger: a bioconductor package for differential expression analysis of digital gene expression data. bioinformatics, 26(1):139–140, 2010.

36. W James Kent, Charles W Sugnet, Terrence S Furey, Krishna M Roskin, Tom H Pringle, Alan M Zahler, and David Haussler. The human genome browser at UCSC. Genome Research, 12(6):996–1006, 2002.

37. Heng Li, Bob Handsaker, Alec Wysoker, Tim Fennell, Jue Ruan, Nils Homer, Gabor Marth, Goncalo Abecasis, and Richard Durbin. The sequence alignment/map format and SAMtools. Bioinformatics, 25(16):2078–2079, 2009.

38. Fidel Ramírez, Friederike Dündar, Sarah Diehl, Björn A Grüning, and Thomas Manke. deepTools: a flexible platform for exploring deepsequencing data. Nucleic Acids Research, 42(W1):W187–W191, 2014.

39. EA Feingold and L Pachter. The ENCODE (ENCyclopedia of DNA elements) project. Science, 306(5696):636–640, 2004.

40. Jason Ernst and Manolis Kellis. ChromHMM: automating chromatin-state discovery and characterization. Nature Methods, 9(3):215–216, 2012.

41. Aaron R Quinlan and Ira M Hall. BEDTools: a flexible suite of utilities for comparing genomic features. Bioinformatics, 26(6):841–842, 2010.

42. Furqan M Fazal, Shuo Han, Kevin R Parker, Pornchai Kaewsapsak, Jin Xu, Alistair N Boettiger, Howard Y Chang, and Alice Y Ting. Atlas of subcellular RNA localization revealed by APEX-Seq. Cell, 178(2):473–490, 2019.

43. Anthony M Bolger, Marc Lohse, and Bjoern Usadel. Trimmomatic: a flexible trimmer for Illumina sequence data. Bioinformatics, 30(15):2114–2120, 2014.

44. Ronny Lorenz, Stephan H Bernhart, Christian Höner zu Siederdissen, Hakim Tafer, Christoph Flamm, Peter F Stadler, and Ivo L Hofacker. ViennaRNA Package 2.0. Algorithms for Molecular Biology, 6(1):1–14, 2011.

45. Zhangming Yan, Norman Huang, Weixin Wu, Weizhong Chen, Yiqun Jiang, Jingyao Chen, Xuerui Huang, Xingzhao Wen, Jie Xu, Qiushi Jin, Kang Zhang, Zhen Chen, Shu Chien, and Sheng Zhong. Genomewide colocalization of RNA–DNA interactions and fusion RNA pairs. Proceedings of the National Academy of Sciences, 116(8):3328–3337, 2019.

46. Zhaokui Cai, Changchang Cao, Lei Ji, Rong Ye, D. Wang, Cong Xia, Sui Wang, Zongchang Du, Naijing Hu, Xiaohua Yu, Juan Chen, Lei Wang, Xianguang Yang, Shunmin He, and Yuanchao Xue. RIC-seq for global in situ profiling of RNA–RNA spatial interactions. Nature, 582(7812):432–437, 2020.

47. Zhipeng Lu, Qiangfeng Cliff Zhang, Byron Lee, Ryan A Flynn, Martin A Smith, James T Robinson, Chen Davidovich, Anne R Gooding, Karen J Goodrich, John S Mattick, et al. Rna duplex map in living cells reveals higher-order transcriptome structure. Cell, 165(5):1267–1279, 2016.

48. Jing Gong, D. Shao, Kui Xu, Zhipeng Lu, Zhi John Lu, Yucheng T Yang, and Qiangfeng Cliff Zhang. RISE: a database of rna interactome from sequencing experiments. Nucleic Acids Research, 46(D1):D194–D201, 2018.

49. Charles E Grant, Timothy L Bailey, and William Stafford Noble. FIMO: scanning for occurrences of a given motif. Bioinformatics, 27(7):1017–1018, 2011.

50. Jessime M Kirk, Susan O Kim, Kaoru Inoue, Matthew J Smola, David M Lee, Megan D Schertzer, Joshua S Wooten, Allison R Baker, Daniel Sprague, David W Collins, Christopher R Horning, Shuo Wang, Qidi Chen, Kevin M Weeks, Peter J Mucha, and J Mauro Calabrese. Functional classification of long non-coding RNAs by k-mer content. Nature Genetics, 50(10):1474–1482, 2018.

51. Mathieu Bastian, Sebastien Heymann, and Mathieu Jacomy. Gephi: an open source software for exploring and manipulating networks. Proceedings of the International AAAI Conference on Web and Social Media, 3(1):361–362, 2009.

52. Uku Raudvere, Liis Kolberg, Ivan Kuzmin, Tambet Arak, Priit Adler, Hedi Peterson, and Jaak Vilo. g:Profiler: a web server for functional enrichment analysis and conversions of gene lists (2019 update). Nucleic Acids Research, 47(W1):W191–W198, 2019.

53. Junichi Iwakiri, Kumiko Tanaka, Takeshi Chujo, Tomohiro Yamazaki, Goro Terai, Kiyoshi Asai, and Tetsuro Hirose. Remarkable improvement in detection of readthrough downstream-of-gene transcripts by semi-extractable RNA-sequencing. RNA, 29(2):170–177, 2023.

54. Chyi-Ying A Chen and Ann-Bin Shyu. AU-rich elements: characterization and importance in mRNA degradation. Trends in Biochemical Sciences, 20(11):465–470, 1995.

55. Mireya Plass, Simon H Rasmussen, and Anders Krogh. Highly accessible AU-rich regions in 3^′^ untranslated regions are hotspots for binding of regulatory factors. PLoS Computational Biology, 13(4):e1005460, 2017.

56. Yuen-Yi Tseng, Branden S Moriarity, Wuming Gong, Ryutaro Akiyama, Ashutosh Tiwari, Hiroko Kawakami, Peter Ronning, Brian Reuland, Kacey Guenther, Thomas C Beadnell, Jaclyn Essig, George M Otto, M Gerard O”Sullivan, David A Largaespada, Kathryn L Schwertfeger, York Marahrens, Yasuhiko Kawakami, and Anindya Bagchi. PVT1 dependence in cancer with MYC copy-number increase. Nature, 512(7512):82–86, 2014.

57. Teresa Colombo, Lorenzo Farina, Giuseppe Macino, and Paola Paci. PVT1: a rising star among oncogenic long noncoding RNAs. BioMed Research International, 2015, 2015.

58. Jing Zhao, Peizhun Du, Peng Cui, Yunyun Qin, Cheng”en Hu, Jing Wu, Zhongwen Zhou, Wenhong Zhang, Lunxiu Qin, and Guangjian Huang. LncRNA PVT1 promotes angiogenesis via activating the STAT3/VEGFA axis in gastric cancer. Oncogene, 37(30):4094–4109, 2018.

59. Seung Woo Cho, Jin Xu, Ruping Sun, Maxwell R Mumbach, Ava C Carter, Y Grace Chen, Kathryn E Yost, Jeewon Kim, Jing He, Stephanie A Nevins, Suet-Feung Chin, Carlos Caldas, S John Liu, Max A Horlbeck, Daniel A Lim, Jonathan S Weissman, Christina Curtis, and Howard Y Chang. Promoter of lncRNA gene PVT1 is a tumor-suppressor DNA boundary element. Cell, 173(6):1398–1412, 2018.

60. Debora Traversa, Giorgia Simonetti, Doron Tolomeo, Grazia Visci, Gemma Macchia, Martina Ghetti, Giovanni Martinelli, Lasse S Kristensen, and Clelia Tiziana Storlazzi. Unravelling similarities and differences in the role of circular and linear PVT1 in cancer and human disease. British Journal of Cancer, 126(6):835–850, 2022.

61. Kevin Tabury, Mehri Monavarian, Eduardo Listik, Abigail K Shelton, Alex Seok Choi, Roel Quintens, Rebecca C Arend, Nadine Hempel, C Ryan Miller, Balázs Györrfy, et al. PVT1 is a stress-responsive lncRNA that drives ovarian cancer metastasis and chemoresistance. Life Science Alliance, 5(11), 2022.

62. Corinne Chureau, Sophie Chantalat, Antonio Romito, Angélique Galvani, Laurent Duret, Philip Avner, and Claire Rougeulle. Ftx is a non-coding rna which affects Xist expression and chromatin structure within the X-inactivation center region. Human Molecular Genetics, 20(4):705–718, 2011.

63. Giulia Furlan, Nancy Gutierrez Hernandez, Christophe Huret, Rafael Galupa, Joke Gerarda van Bemmel, Antonio Romito, Edith Heard, Céline Morey, and Claire Rougeulle. The Ftx noncoding locus controls X chromosome inactivation independently of its RNA products. Molecular Cell, 70(3):462–472, 2018.

64. Andrea Cerase, Alexandros Armaos, Christoph Neumayer, Philip Avner, Mitchell Guttman, and Gian Gaetano Tartaglia. Phase separation drives X-chromosome inactivation: a hypothesis. Nature Structural & Molecular Biology, 26(5):331–334, 2019.

65. Davide Cirillo, Mario Blanco, Alexandros Armaos, Andreas Buness, Philip Avner, Mitchell Guttman, Andrea Cerase, and Gian Gaetano Tartaglia. Quantitative predictions of protein interactions with long noncoding RNAs. Nature Methods, 14(1):5–6, 2017.

66. Sandra L Wolin and Lynne E Maquat. Cellular RNA surveillance in health and disease. Science, 366(6467):822–827, 2019.

67. Jerome M Molleston, Leah R Sabin, Ryan H Moy, Sanjay V Menghani, Keiko Rausch, Beth Gordesky-Gold, Kaycie C Hopkins, Rui Zhou, Torben Heick Jensen, Jeremy E Wilusz, and Sara Cherry. A conserved virus-induced cytoplasmic TRAMP-like complex recruits the exosome to target viral rna for degradation. Genes & Development, 30(14):1658–1670, 2016.

68. Kannanganattu V Prasanth, Supriya G Prasanth, Zhenyu Xuan, Stephen Hearn, Susan M Freier, C Frank Bennett, Michael Q Zhang, and David L Spector. Regulating gene expression through RNA nuclear retention. Cell, 123(2):249–263, 2005.

69. Ankur Jain and Ronald D Vale. RNA phase transitions in repeat expansion disorders. Nature, 546(7657):243–247, 2017.

70. Rut Valgardsdottir, Ilaria Chiodi, Manuela Giordano, Antonio Rossi, Silvia Bazzini, Claudia Ghigna, Silvano Riva, and Giuseppe Biamonti. Transcription of Satellite III non-coding RNAs is a general stress response in human cells. Nucleic Aacids Research, 36(2):423–434, 2008.

71. Caroline Jolly, Alexandra Metz, Jérome Govin, Marc Vigneron, Bryan M Turner, Saadi Khochbin, and Claire Vourc”h. Stress-induced transcription of satellite III repeats. The Journal of Cell Biology, 164(1):25–33, 2004.

72. Tetsuya Kawaguchi, Akie Tanigawa, Takao Naganuma, Yasuyuki Ohkawa, Sylvie Souquere, Gerard Pierron, and Tetsuro Hirose. SWI/SNF chromatin-remodeling complexes function in noncoding RNA-dependent assembly of nuclear bodies. Proceedings of the National Academy of Sciences, 112(14):4304–4309, 2015.

73. Masahiro Onoguchi, Chao Zeng, Ayako Matsumaru, and Michiaki Hamada. Binding patterns of RNA-binding proteins to repeat-derived RNA sequences reveal putative functional RNA elements. NAR Genomics and Bioinformatics, 3(3):lqab055, 2021.

74. Chao Zeng, Atsushi Takeda, Kotaro Sekine, Naoki Osato, Tsukasa Fukunaga, and Michiaki Hamada. Bioinformatics approaches for determining the functional impact of repetitive elements on non-coding RNAs. In piRNA: Methods and Protocols, pages 315–340. Springer US, 2022.

75. Daniel R Garalde, Elizabeth A Snell, Daniel Jachimowicz, Botond Sipos, Joseph H Lloyd, Mark Bruce, Nadia Pantic, Tigist Admassu, Phillip James, Anthony Warland, Michael Jordan, Jonah Ciccone, Sabrina Serra, Jemma Keenan, Samuel Martin, Luke McNeill, E Jayne Wallace, Lakmal Jayasinghe, Chris Wright, Javier Blasco, Stephen Young, Denise Brocklebank, Sissel Juul, James Clarke, Andrew J Heron, and Daniel J Turner. Highly parallel direct RNA sequencing on an array of nanopores. Nature Methods, 15(3):201–206, 2018.

76. Gregory G Fuller, Ting Han, Mallory A Freeberg, James J Moresco, Amirhossein Ghanbari Niaki, Nathan P Roach, John R Yates III, Sua Myong, and John K Kim. RNA promotes phase separation of glycolysis enzymes into yeast G bodies in hypoxia. Elife, 9, 2020.

77. Christiane Iserman, Christine Desroches Altamirano, Ceciel Jegers, Ulrike Friedrich, Taraneh Zarin, Anatol W Fritsch, Matthaus Mittasch, Antonio Domingues, Lena Hersemann, Marcus Jahnel, Doris Richter, Ulf-Peter Guenther, Matthias W Hentze, Alan M. Moses, Anthony A Hyman, Günter Kramer, Moritz Kreysing, Titus M Franzmann, and Simon Alberti. Condensation of Ded1p promotes a translational switch from housekeeping to stress protein production. Cell, 181(4):818–831, 2020.

78. Rena Onoguchi-Mizutani and Nobuyoshi Akimitsu. Long noncoding RNA and phase separation in cellular stress response. The Journal of Biochemistry, 171(3):269–276, 2022.

79. Ryan J Ries, Sara Zaccara, Pierre Klein, Anthony Olarerin-George, Sim Namkoong, Brian F Pickering, Deepak P Patil, Hojoong Kwak, Jun Hee Lee, and Samie R Jaffrey. m6A enhances the phase separation potential of mRNA. Nature, 571(7765):424–428, 2019.

80. Aleksej Drino and Matthias R Schaefer. RNAs, phase separation, and membrane-less organelles: Are post-transcriptional modifications modulating organelle dynamics? BioEssays, 40(12):1800085, 2018.

81. Hong Gil Lee, Jiwoo Kim, and Pil Joon Seo. N6-methyladenosine– modified RNA acts as a molecular glue that drives liquid–liquid phase separation in plants. Plant Signaling & Behavior, 17(1):2079308, 2022.

82. Michael Kertesz, Yue Wan, Elad Mazor, John L Rinn, Robert C Nutter, Howard Y Chang, and Eran Segal. Genome-wide measurement of RNA secondary structure in yeast. Nature, 467(7311):103–107, 2010.

83. Eric L Van Nostrand, Gabriel A Pratt, Alexander A Shishkin, Chelsea Gelboin-Burkhart, Mark Y Fang, Balaji Sundararaman, Steven M Blue, Thai B Nguyen, Christine Surka, Keri Elkins, Rebecca Stanton, Frank Rigo, Mitchell Guttman, and Gene W Yeo. Robust transcriptome-wide discovery of RNA-binding protein binding sites with enhanced CLIP (eCLIP). Nature Methods, 13(6):508–514, 2016.

84. Alessandro Bonetti, Federico Agostini, Ana Maria Suzuki, Kosuke Hashimoto, Giovanni Pascarella, Juliette Gimenez, Leonie Roos, Alex J Nash, Marco Ghilotti, Christopher JF Cameron, Matthew Valentine, Yulia A Medvedeva, Shuhei Noguchi, Eneritz Agirre, Kaori Kashi, Samudyata, Joachim Luginbühl, Riccardo Cazzoli, Saumya Agrawal, M Nicholas Luscombe, Mathieu Blanchette, Takeya Kasukawa, Michiel de Hoon, Erik Arner, Boris Lenhard, Charles Plessy, Gonçalo Castelo-Branco, Valerio Orlando, and Piero Carninci. RADICL-seq identifies general and cell type–specific principles of genome-wide RNA-chromatin interactions. Nature Communications, 11(1):1–14, 2020.

